# Distinct cortical correlates of perception and motor function in balance control

**DOI:** 10.1101/2023.08.22.554282

**Authors:** Jasmine L. Mirdamadi, Lena H. Ting, Michael R. Borich

## Abstract

Fluctuations in brain state alter how we perceive our body and generate movements but have not been investigated in functional whole-body behaviors. During reactive balance control, we recently showed that evoked brain activity is associated with balance ability in healthy young individuals. Further, in individuals with Parkinson’s disease, impairments in whole-body motion perception in reactive balance are associated with clinical balance impairment. Here we investigated brain activity during whole-body motion perception in reactive balance in healthy young adults. We hypothesized that flexibility in brain states underlies successful perception and movement during whole-body movement. We characterized two cortical sensorimotor signals using electroencephalography localized to the supplementary motor area: 1) the “N1 response”, a perturbation-evoked potential that decreases in amplitude with expectancy and is larger in individuals with lower balance function; and 2) pre-perturbation beta oscillatory activity, a rhythm that favors maintenance of the current sensorimotor state and is inversely associated with perception in seated somatosensory perceptual tasks. In a two-alternative forced choice task, participants judged whether pairs of backward support-surface perturbations during standing were in the “same” or “different” direction. As expected, lower whole-body perception was associated with lower balance ability. Within a perturbation pair, N1 attenuation was larger on correctly perceived trials and associated with better balance, but not perception. In contrast, pre-perturbation beta power was higher on incorrectly perceived trials and associated with poorer perception, but not balance. Taken together, flexibility in different cortical processes influences perceptual accuracy but have distinct associations with balance and perceptual ability.

## Introduction

Brain states are known to modulate perception during simple unimodal sensory detection tasks (Jones et al., 2010; Shin et al., 2017) and discrete upper limb movements (Hussain et al., 2022; Little et al., 2019). How brain states modulate complex multisensory processes required for whole body sensorimotor control is unclear. Reactive balance control requires both multisensory integration as well as multi-joint coordination. When balance is perturbed, visual, somatosensory and vestibular inputs are integrated to generate balance-correcting muscle activity (Horak, 2006; Horak and Macpherson, 2011; Welch and Ting, 2008). Sensory integration processes also give rise to conscious perception of the direction of whole-body motion which varies across individuals (Puntkattalee et al., 2016) and could contribute to sensorimotor impairments in individuals with neurologic conditions like Parkinson’s disease (Bong et al., 2020; Halperin et al., 2020). During perceptual tasks in a seated rest position, variations in cortical state prior to a tactile stimulus influences perceptual accuracy (Jones et al., 2010; Shin et al., 2017). In reactive balance, variations in perturbation-evoked activity during balance recovery reflects individual differences in balance ability (Ghosn et al., 2020; Payne and Ting, 2020a). However, the role of pre-perturbation cortical state and perturbation-evoked cortical activity during whole-body motion perception of reactive balance is unknown.

Assessing cortical activity in reactive balance offers a model to examine relationships between perception, movement, and brain states in a functional whole-body context. Even prior to conscious perception of whole-body motion elicited in reactive balance, there is a robust negative event-related potential (N1) 100-200 msec post-perturbation localized to the supplementary motor area (SMA) (Marlin et al., 2014; Mierau et al., 2015; Solis-Escalante et al., 2020). Although the SMA is traditionally thought to contribute to balance through error assessment and motor planning (Nachev et al., 2008; Solis-Escalante et al., 2020; Tanji, 1994), it also receives sensory inputs (Jürgens, 1984; Wiesendanger et al., 1985) and therefore may contribute to perception. N1 responses decrease with prior sensory stimulation (Staines et al., 2001), experience (Mierau et al., 2015; Payne et al., 2019a), and predictability (Adkin et al., 2006), and may contribute to preparing the body for movement (Payne and Ting, 2020b). Thus, N1 modulation may reflect flexibility in preparing sensorimotor responses for perception and movement.

Fluctuations in cortical states modulate perception and movement in nonmobile settings (Hussain et al., 2022; Jones et al., 2010; Little et al., 2019; Shin et al., 2017), but have not been investigated in whole-body behaviors. Beta power is thought to support maintenance of the ‘status quo’ (Engel and Fries, 2010). In the motor domain, beta power is higher at rest and decreases just prior to movement, presumably to release the current sensorimotor state and prepare the system for upcoming sensorimotor processing (Barone and Rossiter, 2021; Kilavik et al., 2013). In the perceptual domain, higher beta power is associated with lower rates of tactile detection in the upper limb (Jones et al., 2010; Shin et al., 2017). Since higher beta power favors maintenance of the current state, lower beta power would indicate greater flexibility and would presumably be associated with better perception and movement.

Here we hypothesized that flexibility in cortical states influences sensory information processing necessary for successful perception and movement. We characterized cortical states localized to the SMA during directional perception of whole-body motion as a function of perceptual accuracy and individual differences in ability. Participants judged whether a pair of standing balance perturbations were in the “same” or “different” direction. We tested whether N1 modulation within a perturbation pair would be larger, reflecting greater flexibility, for trials that are correctly perceived and in individuals with better balance ability. We further tested whether pre-stimulus beta power would be lower, reflecting more flexibility, for correctly perceived trials and in individuals with better perceptual ability. Our findings reveal that perception and balance have distinct neural correlates. Flexibility of perturbation-evoked activity was associated with better balance, whereas flexibility of pre-perturbation state was associated with better perception.

## Materials and Methods

### Participants

19 neurotypical younger adults (24 ± 5 years, 9 female) participated in a single experimental testing session. The experimental protocol was approved by the Emory University Institutional Review board, and all participants gave written informed consent before entering the study.

### Whole-Body Motion Perception

Participants stood barefoot with feet hip-width apart on a moveable platform. Pieces of tape were added to the platform for consistent foot placement and monitored online by the experimenter. Participants wore an overhead harness for safety, stood with their eyes closed, and wore headphones playing white noise to eliminate visual and auditory feedback.

Participants underwent standing balance support-surface translation perturbations where they judged whether a pair of perturbations were in the same or different direction. The first perturbation was always straight backwards, and the second perturbation was backwards with a lateral component (Δθ). Shortly after the end of the second perturbation (< 3 seconds), participants gave a verbal response of “same” or “different” (Figure 1A) (Bong et al., 2020; Puntkattalee et al., 2016). There were two main blocks of perceptual testing.

**Figure 1.**
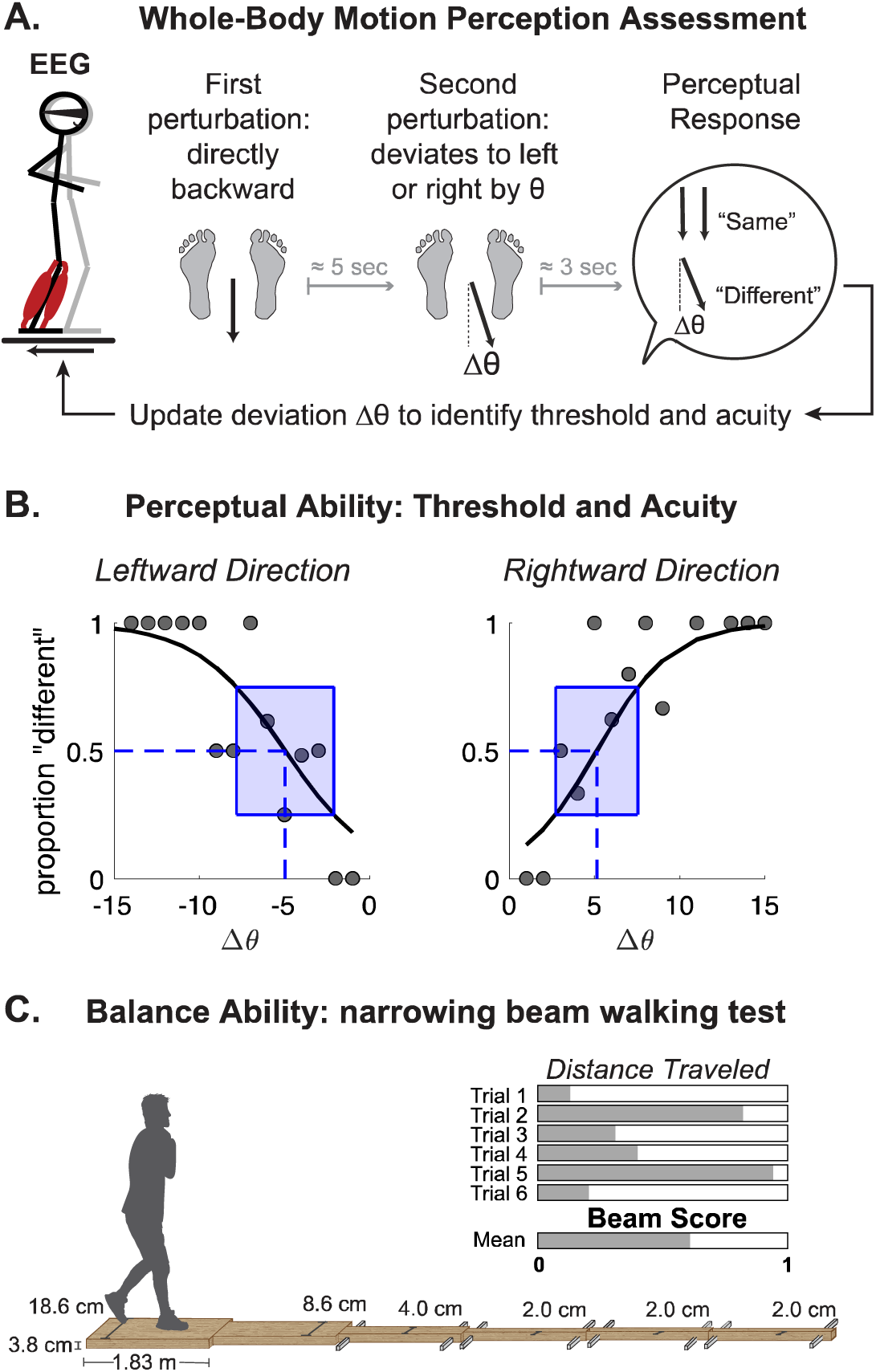
Perception and balance assessments A) Whole-body motion perception protocol. EEG was continuously recorded during pairs of standing balance perturbations. After each pair, participants provided verbal response of whether the perturbations were in the “same” or “different” directions. The second perturbation was identical in amplitude to the first but deviated at an angle that was determined online using an adaptive algorithm to optimize estimation of perceptual acuity and threshold. B) Exemplar perceptual responses for a single participant for perturbation angles in the leftward and rightward direction. Perceptual responses (gray dots) were fit with a logistic function to extract acuity (width of blue shaded rectangle) and threshold (blue vertical dashed line). Acuity defined as the difference between 25th and 75th percentile. Threshold defined as angle at which participants were equally likely to report “same” or “different”. C) Narrowing beam walking test was used as a metric of individual balance ability that required participants to walk one foot in front of the other with their arms across their chest. Beam score was computed as the average proportional distance traveled across 6 trials.

In the first testing block, Δθ was adapted online using the psi method through the Palamedes toolbox in MATLAB (Prins and Kingdom, 2018). Unlike other adaptive algorithms, like PEST (Parameter Estimation by Sequential Testing) (Taylor and Creelman, 1967) and staircase methods (Leek, 2001), the psi method determines the perturbation angle (Δθ) after every trial, based upon the posterior distribution that contains the range of possible values of the slope and threshold parameters. Psi chooses angles that best predict the threshold, and then chooses additional stimulus intensities well above and below the threshold to best predict the slope. The range of possible perturbation angles was linearly spaced from 1 to 30 degrees, based upon thresholds obtained in our previously published work in younger adults (Puntkattalee et al., 2016). A total of 60 pairs of perturbations were given, with right and left angles interleaved.

In the second testing block, Δθ was individualized to each participants’ perceptual threshold for the right and left sides. 108 total pairs of perturbations were delivered. 2/3 of these trials were administered at threshold (50% accuracy, 36 for each direction), and 1/3 were delivered at suprathreshold angles (75% and 95% accuracy, 9 for each magnitude and direction). The order of the direction (left versus right) and angle (Δθ) were delivered in a pseudorandom order to minimize predictability. Participants were given seated rest breaks for ∼3-5 minutes after every 20 trials, or upon request, to minimize fatigue and adaptation.

Perceptual data from the psi block was fit with a psychometric function for the leftward and rightward directions to characterize perceptual acuity and threshold. Acuity was defined as the interquartile range (Figure 1B, dashed blue rectangle) and threshold was defined as the angle at which perturbations were correctly discriminated as different on half of the trials (Figure 1B, dotted pink line) (Mirdamadi and Block, 2020; Ostry et al., 2010; Wilson et al., 2010).

### Balance Ability

Individual balance ability was assessed using a modified version of the narrowing beam-walking test (Sawers and Hafner, 2018) (Figure 1C). To avoid a ceiling effect, the narrowing beam was extended to include two additional segments of the narrowest width (2 cm) for a total distance of 34 feet. Participants were instructed to walk one foot in front of the other with their arms across their chest wearing standardized footwear. There were no requirements for speed, and performance was assessed based on the average total distance traversed over six trials, normalized to the total beam length (Payne and Ting, 2020b). The distance for a trial was based on the toe of the foot that stepped off the beam, or the distance of the front toe when their arms uncrossed.

### EEG recording and analyses

We recorded cortical activity (sampling frequency 1000Hz, impedance <25kOhm) using a 64-channel active electrode cap (actiCAP) and ActiCHamp amplifier with a 24-bit A/D converter and an online 20 kHz anti-aliasing low-pass filter (Brain Products, GmbH). Fz was the online reference electrode and Fpz was the ground electrode.

EEG data were processed using the EEGLAB toolbox (Delorme and Makeig, 2004). Data were high-pass filtered (cutoff 0.5 Hz, finite impulse response, filter order of 3300) and downsampled to 500 Hz. Bad channels were identified using the *clean_rawdata* plugin (flat channel > 5 seconds, high-frequency noise > 4 SDs, or correlation with nearby channels < 0.6). Bad channels were confirmed with visual inspection and interpolated. Data were then re-referenced to an average reference across all channels. 60 Hz line noise was removed with the Zapline-plus plugin (Klug and Kloosterman, 2022). Data were epoched -2 to 2 seconds around each perturbation, and decomposed into maximally independent components (ICs) using adaptive mixture component analysis algorithm (AMICA) (Palmer et al., 2008). ICs from AMICA were categorized using the ICLabel plugin, an automated algorithm that identifies nonbrain sources (e.g., eye, muscle, and cardiac activity) and brain sources, and confirmed with visual inspection (Pion-Tonachini et al., 2019). Nonbrain sources were removed. Brain sources were mapped onto a standard MNI template and estimated using the DIPFIT plugin. Any ICs located outside of the brain or with high residual variance (>15%) from the scalp projection of the best-fitting equivalent dipole were excluded prior to further analyses (Klug and Gramann, 2021; Oostenveld and Oostendorp, 2002). Brain ICs from each participant were clustered by a K-means clustering algorithm in the STUDY portion of EEGLAB using dipole location.

Analyses were performed on the cluster that gave rise to the largest cortical N1 which was localized to the supplementary motor area (SMA) (Figure 4A). If multiple ICs from a single participant clustered together, the IC with the largest percentage power accounted for (ppaf) accounted for between 100-200 post-perturbation was used in analyses (Figure 4C, exemplar IC time course and associated ppaf).

**Figure 4.**
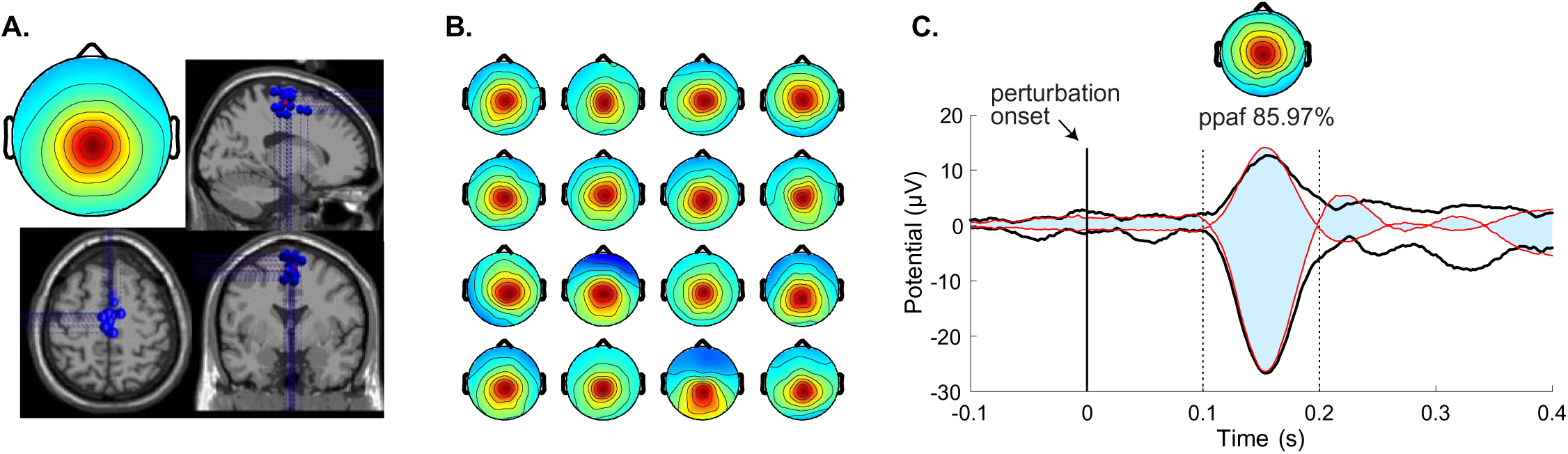
Source-localization of N1 responses. A. Group scalp topography and localization. Centroid of individual ICs (blue dots) localized to Right BA6 Talairach coordinates: (X = 1, Y = -6, Z = 54). B. Scalp maps for each participant showing the topographic distribution of the IC with the largest contribution to the N1 projected across all channels. Red color indicates higher activity over mid-central electrodes. C. IC time series for exemplar subject, with the N1 contributing to 86% power accounted for 100-200 msec post-perturbation (dashed vertical lines). Thick black lines denote data envelope – the maximum and minimum activation of all brain ICs projected across all electrodes. Red line denotes envelope for IC that contributed largest to the N1 response. Blue shaded area denotes proportion of signal

To assess cortical responses post-perturbation, we extracted single trial N1 amplitudes that were baseline subtracted (-150 to -50 msec before perturbation onset) for each participant. The N1 was quantified as the largest negative peak 100-200 msec after perturbation onset.

To assess cortical states prior to the perturbation, we extracted condition mean and single trial pre-perturbation power spectra from -1000 to 0 msec using Welch’s method (nonoverlapping hamming windows, 500-window length) for each participant. The FOOOF (Fitting Oscillations & One-Over-F) toolbox (Donoghue et al., 2020) was used to decompose power spectra into aperiodic (1/f) and periodic components (activity above 1/f) from 2 to 35 Hz using the parameters: minimum peak height: 0.1, minimum peak width: 2, max number of peaks: 4, peak threshold: 2, knee parameter: ‘fixed’. Peak periodic beta power was extracted from the fitted spectra from 13-30 Hz (Figure 2B, pink vertical dashed line). If more than one peak was detected from 13-30 Hz, peak beta power was averaged across the peaks. Since the width of peaks could also vary (Figure 2B – left versus right column), we further extracted area under the fooofed spectra between 13-30 Hz (Figure 2B, pink shared area). Two representative example participants with contrasting perceptual thresholds had distinct differences in the magnitude of pre-stimulus beta power (Figure 2, left column versus right column). The individual with a lower perceptual threshold (Figure 2, left column) had lower pre-stimulus beta power for the nonparameterized power spectra (Figure 2A) and the parameterized periodic beta power (Figure 2B, pink dashed line and pink shared area) compared to the individual with a higher perceptual threshold (Figure 2, right column).

**Figure 2.**
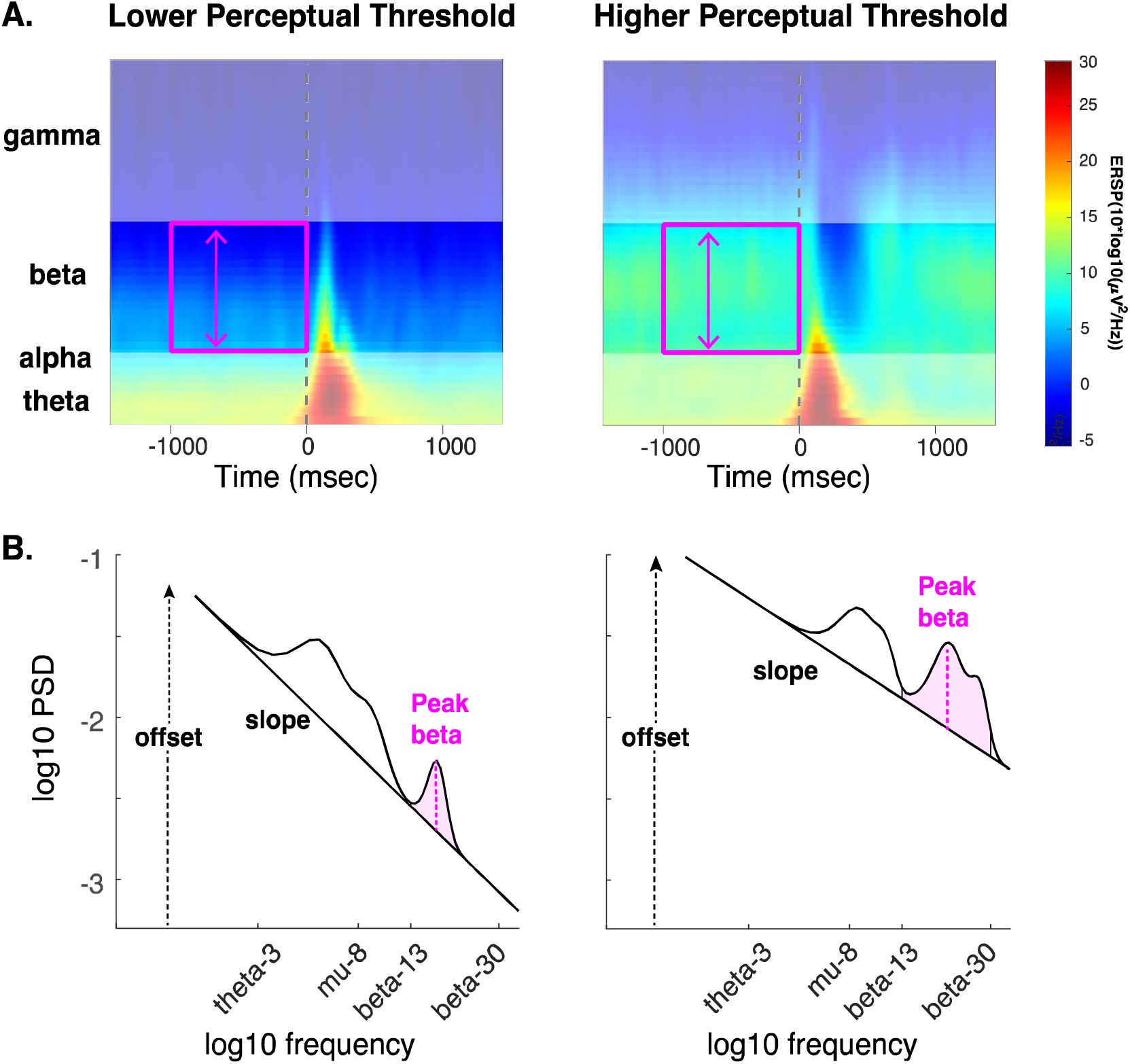
Exemplar spectral data from two participants with different perceptual abilities. A. time-frequency spectrograms pre-and post-perturbation. Pink rectangle denotes beta range (13-30Hz) analyzed pre-perturbation (-1000 to 0 msec). B. log-transformed pre-stimulus (-1000 – 0 msec) power spectral density parameterized into periodic and aperiodic components. Aperiodic power (1/f component that is linear with log transformation) described by the slope and offset. Periodic power represented by peaks above the aperiodic component. Beta power was described using peak power (pink dashed line) and area under the curve (pink shaded region) from 13-30 Hz in the 1000 msec prior to the perturbation. Individual with lower perceptual threshold (better ability) had lower pre-stimulus beta power (left column) compared to the individual with higher perceptual threshold (right column).

### Statistical Analyses

To test the interrelationships between perception, movement, and brain states, we first examined associations between whole-body motion perception and balance ability. We tested whether perceptual ability differed between the left and right sides using paired samples t-tests. We then tested associations between whole-body motion perceptual ability and balance ability using independent correlations. Separate analyses were performed for acuity and threshold.

To examine whether flexibility in brain states affects perception *within* individuals, we ran separate linear mixed effects models on N1 amplitude and pre-stimulus beta power using predictor variables of Perturbation Order and Accuracy using the lmerTest package in R using the following formula:

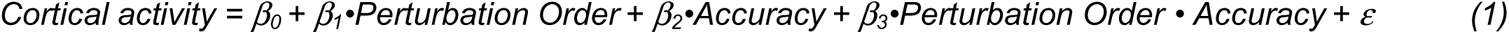

Perturbation order and Accuracy served as fixed effects, with reference levels of First (vs. Second) and Correct (vs. Incorrect). Correct trials were those that were reported to be “different”. Participants served as a random effect. *T*o examine whether cortical activity changed overtime with repeated trials, we also ran analyses on single trial EEG data using the above model with the addition of trial number as a categorical variable.

To examine whether flexibility in brain states affects perception and balance *between* individuals, we first ran separate independent correlations between N1 modulation (ΔN1 = N1 amplitude second perturbation – first perturbation) and pre-stimulus beta power with ability (balance, acuity, threshold). We also performed multiple regression to evaluate balance, acuity, and threshold as predictors of ΔN1 and pre-stimulus beta power.

To confirm the role of oscillatory beta activity in perception and balance rather than broadband 1/f activity, we ran parallel analyses (both within individuals and between individuals) on the aperiodic offset and aperiodic slope.

Since the psi block was adaptive, the angles ranged from 0 to 30 degrees depending on each individual’s response and therefore variable across individuals. This block was used for quantifying each individual’s perception (Figure 1B, acuity and threshold). All statistical analyses on cortical activity were performed on the individualized block since this block had the largest number of trials that was at an equivalent level of perceptual challenge (i.e., threshold) across all participants. Two out of the 19 participants only had the psi block and were excluded from cortical activity analyses.

## Results

### Perception-Balance Associations

Perceptual acuity, the range of angles for which participants were correct in 25 to 75% of trials (Figure 3A, shaded rectangles) ranged between 2.67 and 12.35° (5.32 ± 2.59°), and perceptual threshold, the angle at which participants were correct 50% of the time (Figure 3A, vertical dashed lines) ranged between 2.59 and 12.77° (7.9 ± 2.73°). These values represent averages across left and right directions, which were not significantly different (Acuity: left – 4.6 ± 2°, right – 6.2 ± 4° [mean ± SD], p = 0.11; Threshold: left – 7.6 ± 3°, right – 8.2 ± 3°, p = 0.43).

**Figure 3.**
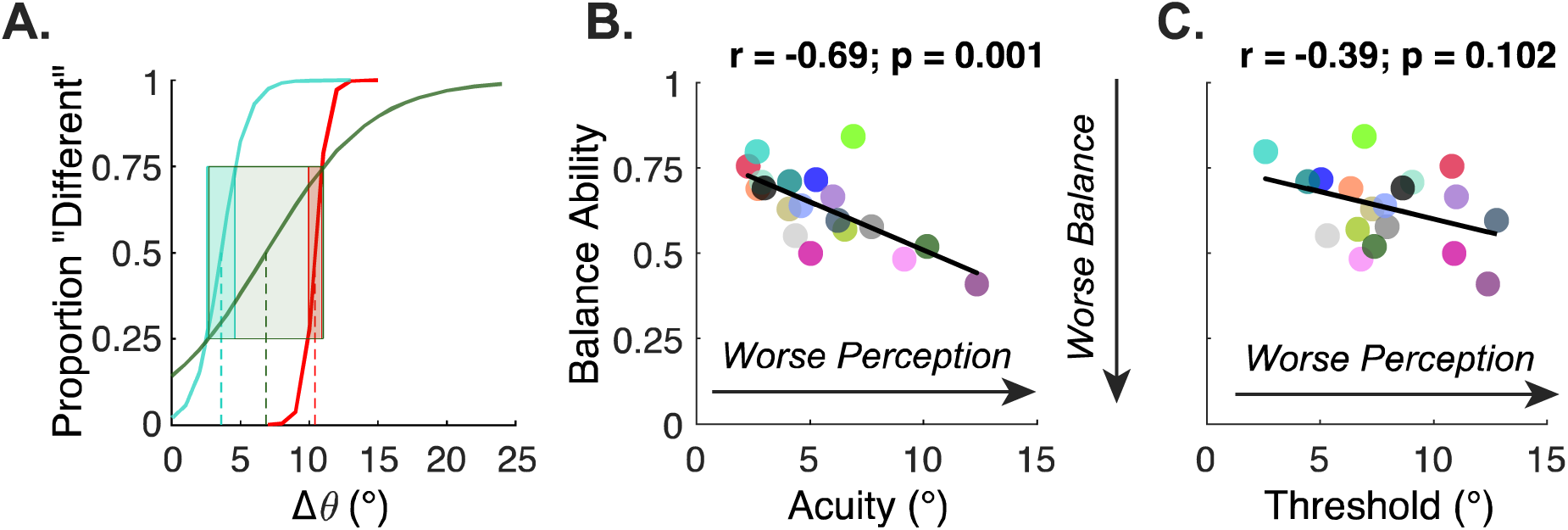
Individual differences in perception and balance ability. A. Exemplar psychometric curves during whole-body motion perception task for three participants with varying acuity (shaded rectangles) and threshold (vertical dashed lines). Two participants (turquoise, red) with relatively better acuity (narrower rectangles, smaller interquartile ranges) despite contrasting thresholds (turquoise – better, red – worse) had better balance. Third exemplar participant (green) had better threshold but worse acuity and had relatively worse balance. B. Lower perceptual acuity was associated with lower balance ability. C. Perceptual threshold was not associated with balance ability.

Across individuals, threshold and acuity were not associated (r = 0.29, p = 0.22). While some individuals had a low threshold (better) and high acuity (better) (Figure 3A – turquoise trace), others had high threshold (worse) and high acuity (better) (Figure 3A – red trace).

Individual variations in perceptual acuity but not threshold was associated with balance ability (normalized distance: range= 0.41 to 0.84; 0.63 ± 0.11). As an example, an individual with relatively lower (i.e., better) threshold but worse acuity had relatively lower balance function (Figure 3A – green trace). In contrast, an individual with relatively higher (i.e., worse) threshold but better acuity had better balance function (Figure 3A – red trace). Across individuals, worse whole-body motion perception acuity was associated with lower balance ability (Figure 3B: r = -0.69, p = 0.001). While a similar direction of association was observed for perceptual threshold with balance ability, the correlation was not statistically significant (Figure 3C: r = -0.39, p = 0.102).

### Source localization

Consistent with prior studies (Marlin et al., 2014; Mierau et al., 2015; Solis-Escalante et al., 2020), the source that contributed largest to the N1 was localized to the SMA (Figure 4A, individual equivalent current dipoles denoted by blue circles). IC scalp maps were similar across participants and illustrate a midfrontal topography (Figure 4B). The estimated location, residual variance, and percentage power accounted for (ppaf) the IC that gave rise to the largest N1 potential between 100-200 msec post-perturbation (Figure 4C, exemplar participant) for each participant is shown in Table 1. One participant was excluded due to high electrode impedance from most sensors related to hairstyle.

**Table 1.** Characteristics of primary source contributing to the N1 potential for each participant. Talairach coordinates of the IC that contributed largest to the N1 potential 100-200 msec post-perturbation. Ppaf - percentage power accounted for. RV – residual variance.

| ID | x | y | z | Ppaf | RV |
| --- | --- | --- | --- | --- | --- |
| 1 | -6.21 | -11.64 | 71.97 | 96.66 | 2.35 |
| 2 | 10.06 | -8.69 | 67.68 | 85.93 | 4.92 |
| 3 | -0.42 | -13.69 | 63.13 | 80.15 | 2.49 |
| 4 | 4.16 | 1.42 | 54.19 | 85.97 | 2.85 |
| 5 | 2.64 | -9.44 | 61.52 | 65.81 | 4.28 |
| 6 | -0.98 | -19.1 | 57.47 | 87.85 | 1.35 |
| 7 | -1.24 | -8.15 | 72.1 | 79.27 | 3.97 |
| 8 | -0.29 | -25.18 | 55.93 | 79.16 | 2.46 |
| 9 | 3.31 | 8.8 | 52.48 | 74.11 | 1.46 |
| 10 | 0.29 | -15.51 | 68.45 | 96.49 | 0.77 |
| 11 | -5.99 | -20.13 | 50.21 | 79.93 | 1.6 |
| 12 | 2.18 | -9.27 | 57.93 | 92.48 | 1.52 |
| 13 | -4.09 | -20.95 | 73.22 | 87.57 | 1.43 |
| 14 | 2.26 | -10.52 | 64.54 | 74.7 | 1.12 |
| 15 | -5.57 | -1.26 | 47.39 | 62.78 | 2.89 |
| 16 | 2.98 | -13.05 | 48.85 | 93.71 | 2.75 |
| mean | 0.19 | -11.02 | 60.44 | 82.66 | 2.39 |

### Task- and individual-specific perturbation-evoked N1 responses

N1 amplitude on a single perturbation was smaller on the second compared to the first and decreased across trials for both correct (i.e., “different”) and incorrect responses (i.e., “same”). Mean N1 amplitude was smaller on the second perturbation compared to the first (β_Perturbation Order_= -1.99 µV, 95% CI: [-2.32, -1.65], t_45_ = -11.95, p < 0.001) (Figure 5, pink versus black bars). The N1 amplitude on either the first or the second perturbation was similar regardless of perceptual accuracy (p = 0.28) (Figure 5, A versus B). These results were further substantiated using single-trial N1 amplitudes; there was a significant decrease in N1 amplitude within a perturbation pair (β_Perturbation Order_ = -2.04 µV, 95% CI: [-2.24, - 1.84], t_2948.02_ = -20.11, p < 0.001). N1 amplitude also decreased across trials (β_Trial Number_ = -0.01, 95% CI: [-0.01, -0.01], t_2949.51_ = -8.15, p < 0.001) regardless of perceptual accuracy (p = 0.12).

**Figure 5.**
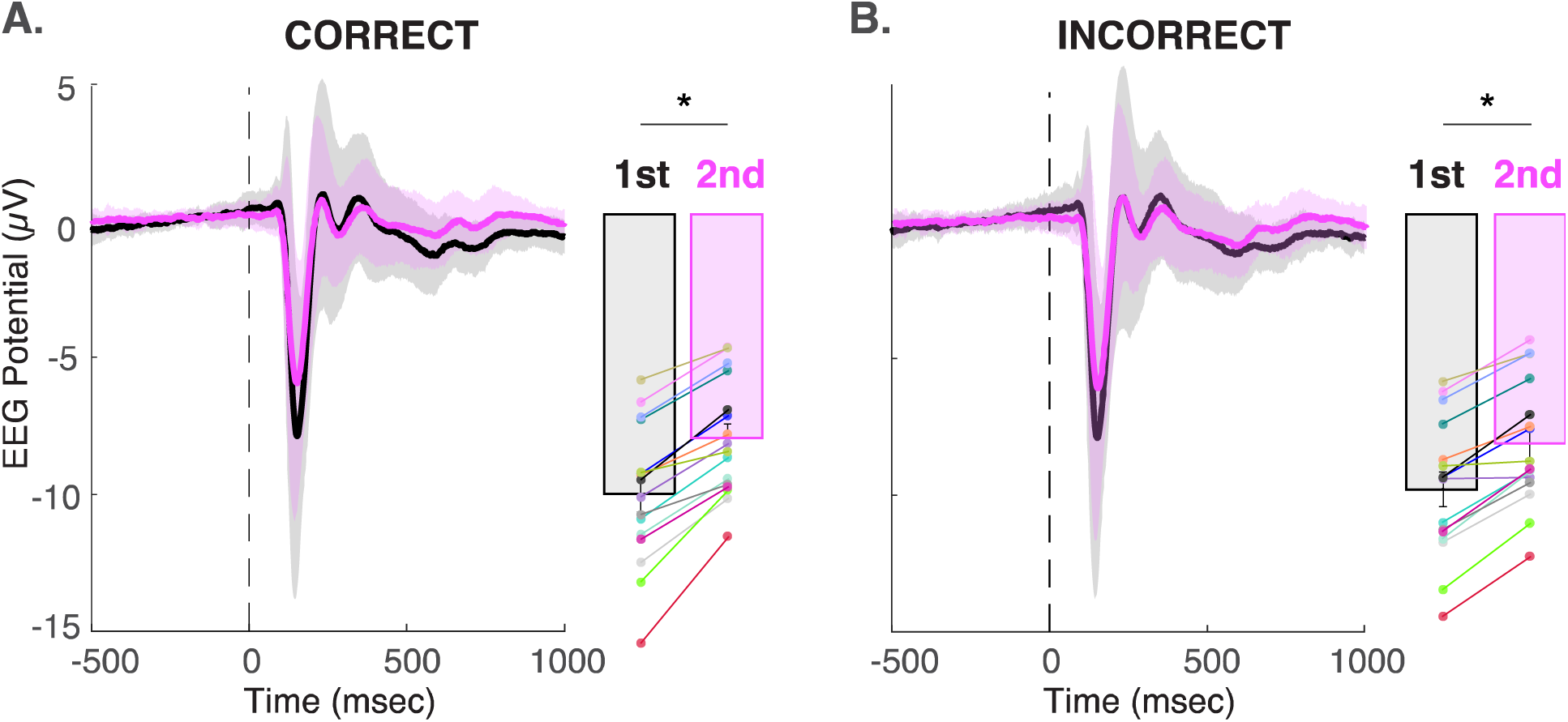
Perturbation-evoked potentials and N1 amplitudes extracted from SMA source across perturbation conditions. Thick solid line and shaded region of ERP trace denote mean and standard deviation, respectively. Bars denote mean N1 amplitude. Error bars denote standard error of the mean. N1 amplitude decreased on the second perturbation relative to the first perturbation for both correctly perceived trials (A.) and incorrectly perceived trials (B.) * Denotes effect of Perturbation Order: p<0.001

Since N1 decreased within a perturbation pair, we ran a parallel analysis on single trial N1 modulation (ΔN1 = N1 amplitude on second – first perturbation). N1 modulation within a perturbation pair was larger on correctly perceived trials versus incorrectly perceived trials (Figure 6A, β_Accuracy_ = 0.42, 95% CI: [0.11, 0.72], t_1528.38_ = 2.65, p = 0.008) and decreased with repeated trials (β_Trial Number_ = -0.01, 95% CI: [- 0.01, -0.00], t_1534.81_ = -2.21, p = 0.028).

**Figure 6.**
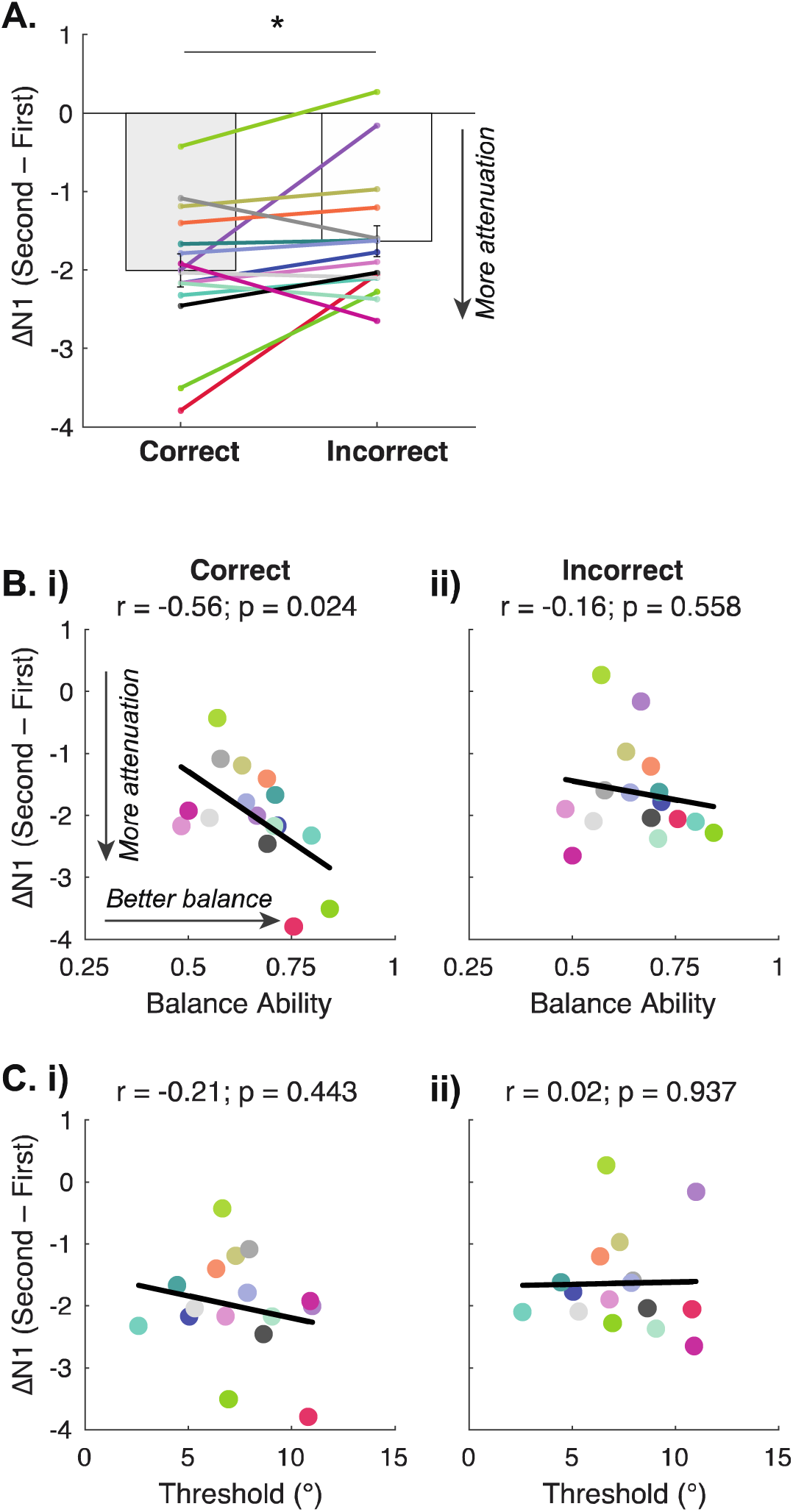
N1 modulation across individuals within a pair of perturbations versus ability. A. N1 attenuation (more negative ΔN1) was larger for correctly perceived trials compared to incorrectly perceived trials. * denotes p=0.008. Error bars denote standard error of the mean. B. Larger N1 attenuation was associated with better balance ability for correctly perceived trials (i) but not incorrectly perceived trials (ii). C. There were no associations between N1 attenuation and perceptual threshold for neither correctly perceived trials (i) nor incorrectly perceived trials (ii).

Across individuals, better balance ability was associated with larger N1 attenuation, but only for the correctly perceived perturbations (correct: r = -0.56, p = 0.02, Figure 6Bi; incorrect: r = -0.16, p = 0.56, Figure 6Bii). Neither perceptual threshold (Figure 6Ci and ii, all p>0.39) nor acuity were associated with N1 attenuation, regardless of perceptual accuracy. The specificity of the associations between balance ability but not perceptual ability on N1 modulation was confirmed with multiple regression, using predictors of balance, threshold, acuity, and accuracy. The multiple regression revealed an interaction between balance and accuracy on N1 modulation (β_Balance x Accuracy_ = 5.09, 95% CI: [1.27, 8.92], t_12_ = -2.90, p = 0.013). The other predictors of acuity, threshold, and any interaction with accuracy on ΔN1 were not significant (all p>0.08). N1 associations with balance ability were specific to N1 attenuation within a pair; There were no associations between N1 amplitude evoked by either the first or second perturbation and ability (all p> 0.19).

### Task- and individual specific pre-perturbation cortical state

Pre-perturbation peak beta power was higher when perception was incorrect compared to when perception was correct (β_Accuracy_ = 0.07, 95% CI: [0.015, 0.125], t_45_ = 2.55, p = 0.014) but did not change within a perturbation pair (β_Perturbation Order_ = 0.02, 95% CI: [-0.032, 0.078], t_45_ = 0.85, p = 0.40) (Figure 7A). This pattern was also observed for beta AUC (β_Accuracy_= 0.81, 95% CI: [0.12, 1.50], t_45_ = 2.36, p = 0.023; β_Perturbation Order_ = -0.14, 95% CI: [-0.83, 0.55], t_45_ = -0.42, p = 0.68) (Figure 7B).

**Figure 7.**
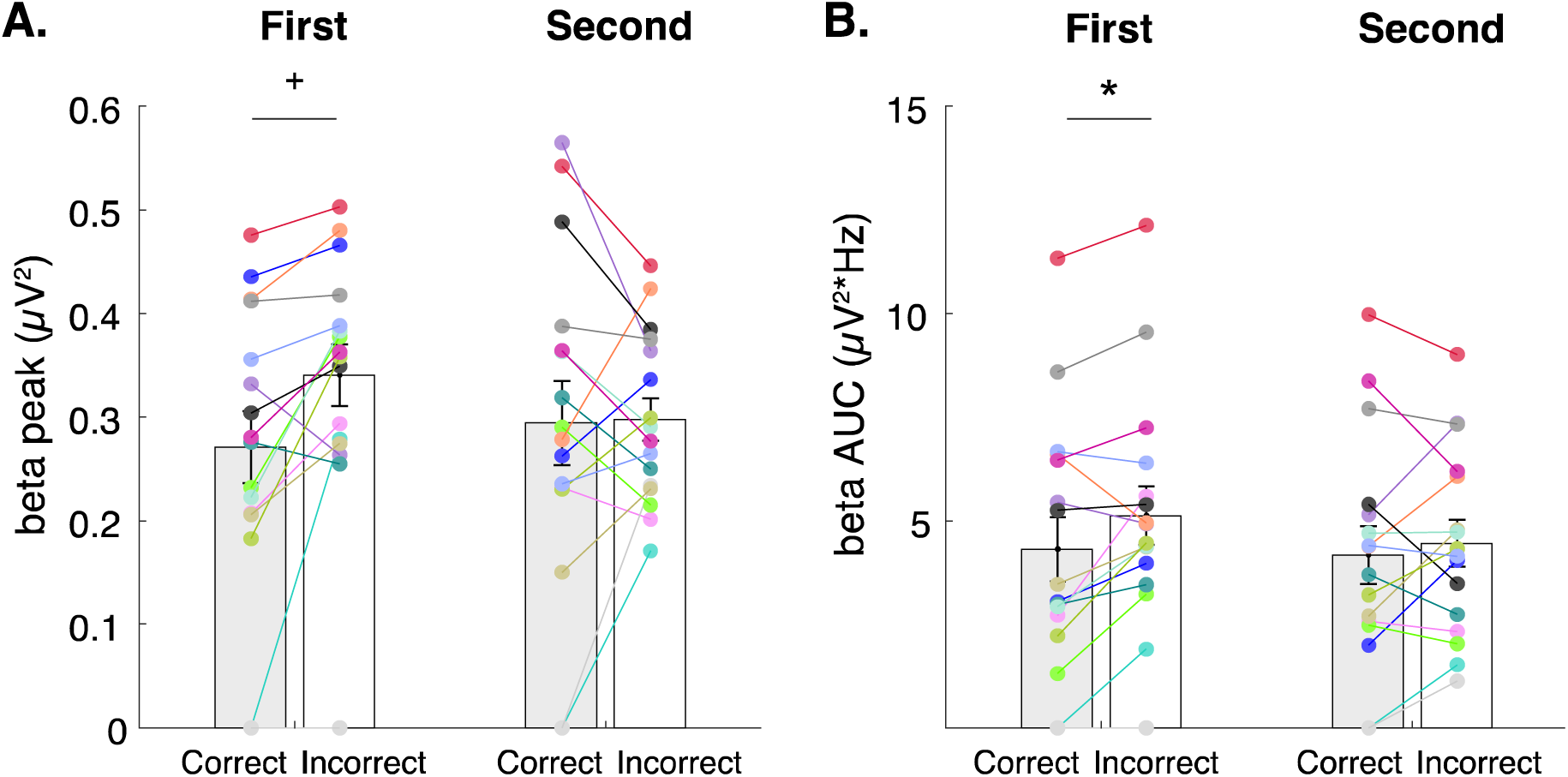
Pre-stimulus beta activity across perturbation conditions. Pre-stimulus beta power was higher when perception was incorrect compared to correct, but only on the first perturbation of the pair. There were no differences in beta power with accuracy for the second perturbations of the pair. There was a similar pattern for peak beta power (A) and beta area under the curve (AUC) (B). + denotes p = 0.014; * denotes p = 0.023. Error bars denote standard error of the mean.

Single trial pre-perturbation beta power analyses revealed that beta power was higher for incorrectly perceived trials compared to correctly perceived trials, but only for the first perturbation (β_Accuracy x Perturbation Order_ = -0.08, 95% CI: [-0.13, -0.02], t_2879.72_ = -2.69, p = 0.007; β_Accuracy_ = 0.05, 95% CI: [0.01 – 0.09], t_2886.31_ = 2.32, p = 0.021). Peak beta power was not systematically modulated with repeated trials (p = 0.71). A similar pattern was observed for single trial beta AUC, with higher beta power on the first compared to the second perturbation but only for incorrect trials (β_Accuracy x Perturbation Order_ = 1.07, 95% CI: [-2.11, -0.03], t_2878.32_ = -2.03, p = 0.043), with no systematic change in power across repeated trials (p = 0.62).

There was no effect of perceptual accuracy on pre-perturbation aperiodic parameters (exponent, offset). Both the exponent and offset showed a small but significant increase within a perturbation pair (Exponent: β_Perturbation Order_ = 0.05, 95% CI: [0.01, 0.10], t_45_ = 2.42, p = 0.020; Offset: β_Perturbation Order_ = 0.07, 95% CI: [0.01, 0.12], t_45_ = 2.56, p = 0.014). However, single trial analyses of these parameters revealed no significant effects or interactions.

Across indiviuduals, lower perceptual thresholds (i.e., better ability) were associated with lower pre-stimulus beta AUC, regardless of perceptual accuracy (Fig 8Bi, ii) (correct: r = 0.76, p = 0.001; incorrect: r = 0.73, p = 0.001). Neither perceptual acuity nor balance ability (Figure 8Ai, ii) were associated with pre-stimulus beta power. These results were confirmed with multiple regression using predictors of balance, threshold, acuity, and accuracy, on pre-stimulus beta power. The multiple regression revealed that only threshold predicted pre-stimulus beta power (β_Threshold_ = = 0.92, 95% CI: [0.45,1.39], t_13.6_ = 4.18, p = 0.001). The other predictors of acuity, balance, and any interaction with accuracy were not significant (all p>0.16).

**Figure 8.**
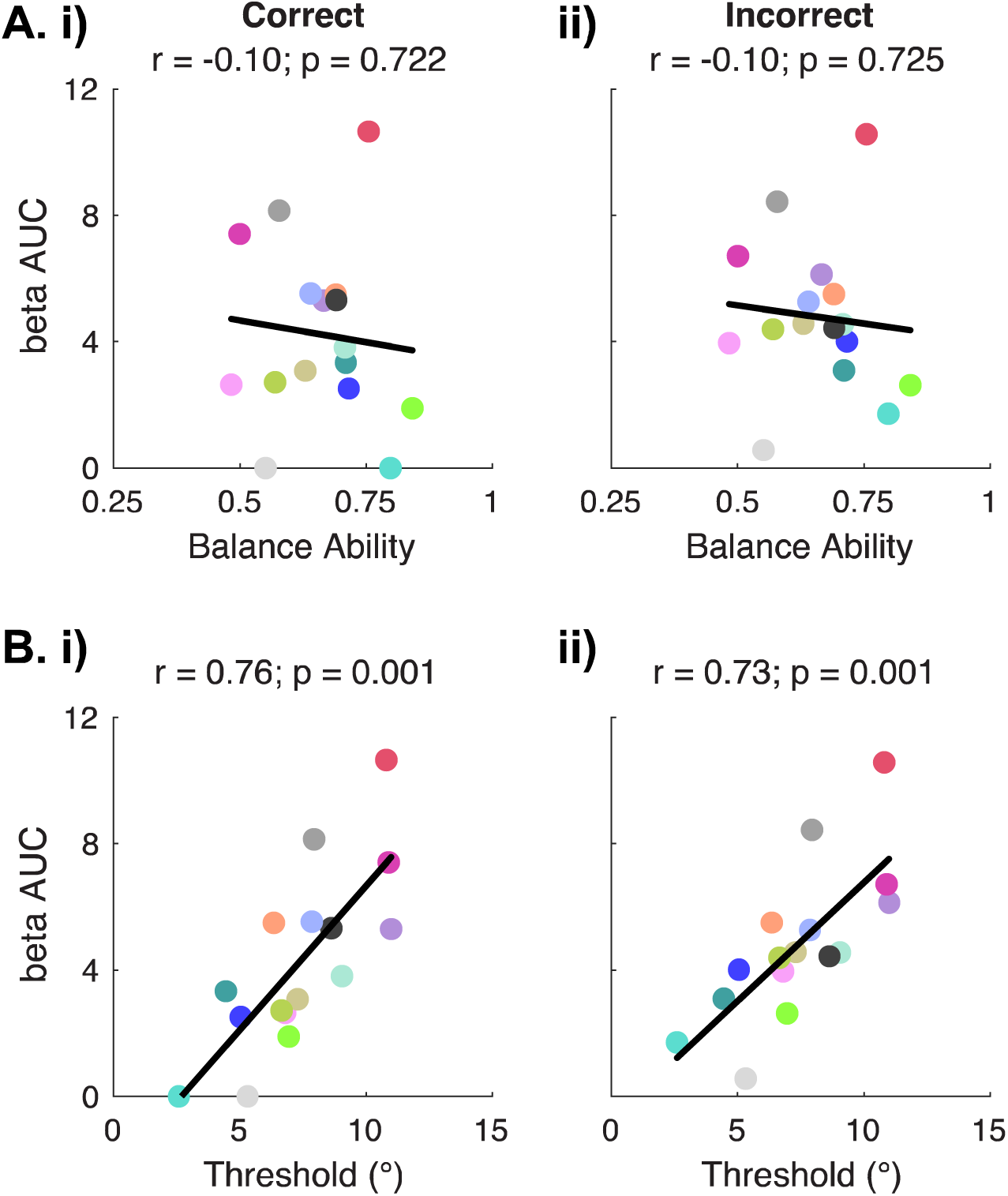
Pre-stimulus beta power across individuals versus ability. A. Pre-stimulus beta power was not associated with balance ability for neither correctly perceived trials (i) nor incorrectly perceived trials (ii). B. Individuals with higher pre-stimulus beta power had higher (i.e., worse) thresholds for both correctly perceived trials (i) and incorrectly perceived trials (ii).

There were no associations between aperiodic parameters and acuity, threshold, or balance (all p>0.11 for independent correlations), supporting the specific role of periodic oscillatory beta power in explaining individual differences in threshold rather than a combination of oscillations and broadband power.

## Discussion

Here we show for the first time, relationships between brain states, perception, and movement that contribute to functional balance control. We identified two distinct cortical activity metrics of flexibility that vary across trials with perceptual accuracy and are uniquely associated with individual differences in balance and perceptual ability. First, we propose that attenuation of balance-perturbation evoked N1 potentials reflects cortical flexibility important for balance control. Attenuation of cortical N1 potentials within a pair of perturbations during a whole-body perceptual task was larger for correctly perceived trials and across individuals was associated with better balance ability. This attenuated brain activity may reflect the ability of the nervous system to adapt its response to predictable future events. Second, we propose that higher beta power reflects a maintenance of the sensorimotor status quo, reducing the system’s flexibility to processing sensory inputs. Beta power was higher on incorrectly perceived trials and across individuals was associated with higher (i.e., worse) perceptual thresholds. Beta power may thus reflect cortical mechanisms that alter sensory processing necessary for perception. These linkages between brain and behavior provide insight into cortical mechanisms underlying individual differences in sensorimotor function that may be important in health and disease.

### Interactions between perceptual and balance ability

The positive association between directional perception and balance ability highlights the important role of multisensory integration processes in maintaining balance. A similar relationship was previously found between impaired directional perception and clinical balance function using the miniBEST in Parkinson’s disease (PD), but not healthy older adults, potentially due to a ceiling effect of the miniBEST in the older adults (Bong et al., 2020). Here, we used the narrowing beam walking task as a more sensitive tool to detect individual differences in balance function (Sawers and Hafner, 2018), revealing associations between whole-body perceptual acuity and balance ability in younger adults. Surprisingly, there was no association between threshold and balance ability, highlighting the importance of characterizing both acuity and threshold of psychometric functions. Perceptual acuity and threshold are not necessarily dependent (Hoseini et al., 2015); an individual can be highly sensitive but have a large threshold, while another individual can be highly sensitive and have a small threshold. We speculate that higher acuity reflects higher certainty of perception, resulting in more efficient adjustments to subtle changes in posture for successful balance. The lack of association between balance and threshold may suggest that individuals compensate for any bias in perceived body position (i.e., threshold) using vision while walking across a beam.

### Attenuation of perturbation evoked N1 potentials reflects cortical flexibility important for balance control

The N1 was localized to SMA, an area that serves as a hub for integrating sensory, motor, and cognitive information to rapidly evaluate and respond to errors (Bonini et al., 2014; Nachev et al., 2008; Tanji, 1994). Evaluating an error signal involves a comparison of expected versus actual sensory feedback which presumably could affect directional perception. However, we observed similar N1 amplitudes regardless of the perceptual response, suggesting a distinction between initial error processing and perception. Decreases in N1 amplitude within a perturbation pair and across trials is consistent with prior literature suggesting that the N1 is influenced by perturbation predictability and experience (Adkin et al., 2006; Mierau et al., 2015; Payne et al., 2019a), potentially mediated by cognitive control (Payne et al., 2019b).

N1 attenuation within a perturbation pair may reflect cortical flexibility for processing predictable sensory input that varies within and between individuals. Because the second perturbation in each pair occurred at predictable timing and magnitude, individuals could anticipate and prepare for the perturbation and adapt with repeated trials, resulting in smaller N1 amplitudes on the second perturbation. Greater N1 attenuation for correctly perceived trials suggests better predictability of the expected postural state thereby enhancing the system’s sensitivity to upcoming sensory inputs. The N1 attenuation observed here is similar to cortical flexibility observed in unimodal paired pulse paradigms where early cortical responses are smaller on the second stimulus relative to the first (Cromwell et al., 2008), thought to be mediated by sensory gating. Sensory gating normally occurs to prevent overflow of sensory information and optimize neural resources for processing novel inputs (Azim and Seki, 2019). Here, greater gating could serve to filter out irrelevant sensory information that increases the salience of the second perturbation to accurately perceive whole-body motion. While prior studies have linked reduced sensory gating to reductions in tactile perception in aging (Cheng and Lin, 2013; Spooner et al., 2019) and neurological dysfunction (Arpin et al., 2017; Nguyen et al., 2020), this is the first study to demonstrate the functional relevance of reduced cortical flexibility to trial-by-trial reductions in whole-body motion perception accuracy.

Between individuals, the attenuation of N1s within a pair may reflect cortical flexibility necessary for successful balance control, since larger N1 attenuation was associated with better balance ability. We speculate that reduced N1 modulation indicates less cortical flexibility that may be inefficient for adapting to subtle changes in sensory input (Azim and Seki, 2019) in balance control. Modulating the N1 may reflect the modulation of motor or cognitive responses with changes in task demands or conditions, or “central set” necessary for successful balance control (Horak et al., 1989). In older adults with and without PD, the inability to change the central set has been linked to inflexibility in postural responses and balance impairments (Chong et al., 2000; Payne et al., 2021). Here, we demonstrate that flexibility in central set also varies across young adults as a function of balance ability. The specificity of the associations between balance ability and N1 modulation rather than absolute N1 amplitude suggests that it is the ability to change sets, i.e., alter responses to predictable sensory input, rather than a single error signal that has functional relevance to balance control. The lack of association with N1 modulation and perceptual ability further points to a distinction between initial sensory processing versus higher-order processes that may guide intrinsic differences in perception.

### Beta oscillations reflect cortical flexibility for whole-body motion perception

Our data suggest that pre-stimulus beta power sets the sensitivity of the perceptual system within and between individuals. Here we show that the inhibitory role of beta processes previously shown in simple unimodal detection or single joint movement tasks in a seated position (Jones et al., 2010; Little et al., 2019; Shin et al., 2017) are also relevant during functional whole-body behaviors. Lower beta power may represent more effective sensory processing, potentially by increasing the salience of somatosensory signals to better discriminate the direction of whole-body motion. In particular, we show that beta fluctuations governing trial-by-trial perception behavior (Little et al., 2019; Shin et al., 2017) are also relevant in a whole-body sensorimotor task. Higher pre-stimulus beta power for incorrectly perceived judgements is consistent with the role of beta is maintaining the current motor state (Engel and Fries, 2010), potentially making the system less receptive to sensory input that would be necessary to correctly discriminate the direction of whole-body motion.

Further support for the inhibitory influence of beta power on perceptual function comes from the finding that individuals with higher pre-stimulus beta power had higher (i.e., worse) perceptual thresholds. If beta power reflects inhibitory processing, lower pre-stimulus beta power may reflect a more permissive state of the system for upcoming somatosensory processing and/or enhance the salience of somatosensory signals (Engel and Fries, 2010; Jones et al., 2010; Shin et al., 2017).

Similarly, individuals exhibiting lower beta power may have more effective sensory processing resulting in lower (i.e., better) discrimination thresholds. Conversely, higher beta power in individuals with worse ability may serve to favor the current sensorimotor state that results in less flexibility (Engel and Fries, 2010), potentially at the expense of processing new sensory information elicited by balance perturbations, and higher discrimination thresholds. Since we parameterized power spectra into periodic and aperiodic components, these findings are unlikely to be driven by non-rhythmic activity, but instead reflect individual differences in beta oscillatory power. The specific role of beta oscillatory power in perception was further substantiated by the findings that neither aperiodic exponent nor offset correlated with perceptual ability.

### Variations in brain state modulate perception and movement

Our findings extend the role of brain states in perception and movement, traditionally studied in isolation during simple tasks, to functional balance control that requires both perception and movement. The distinct associations between flexible brain activity and functional behaviors have important implications for understanding mechanisms of perceptual-motor interactions in people with impaired balance, such as with aging, stroke, and PD. Reduced sensory gating and elevated beta power previously shown with aging, stroke, and PD may reflect different mechanisms of reduced cortical flexibility necessary for perception and movement (Azim and Seki, 2019; Brown, 2003; Espenhahn et al., 2020; Rossiter et al., 2014; Thibaut et al., 2017). A better understanding of brain states during functional balance control may help dissociate balance and perceptual deficits and help guide mechanistic-based interventions for rehabilitation.

## Acknowledgements

This work was supported by National Institutes of Health Eunice Kennedy Shriver National Institutes of Child Health & Human Development (F32HD105458 to JLM), National Institute of Neurological Disorders and Stroke (5T32NS007480-15 to JLM), and National Institute on Aging (R01 AG072756 to LHT and MRB). The authors would like to acknowledge Dr. Sara Hussain for her assistance in spectral parameterization methods, and Scott Boebinger, Kennedy Kerr, and Rish Rashtogi for their assistance with data collection.

## References

Adkin, A.L., Quant, S., Maki, B.E., McIlroy, W.E., 2006. Cortical responses associated with predictable and unpredictable compensatory balance reactions. Experimental Brain Research 172, 85–93. 10.1007/s00221-005-0310-9

Arpin, D.J., Gehringer, J.E., Wilson, T.W., Kurz, M.J., 2017. A reduced somatosensory gating response in individuals with multiple sclerosis is related to walking impairment. Journal of Neurophysiology 118, 2052–2058. 10.1152/jn.00260.2017

Azim, E., Seki, K., 2019. Gain control in the sensorimotor system. Curr Opin Physiol 8, 177–187. 10.1016/j.cophys.2019.03.005

Barone, J., Rossiter, H.E., 2021. Understanding the Role of Sensorimotor Beta Oscillations. Front Syst Neurosci 15, 655886. 10.3389/fnsys.2021.655886

Bong, S.M., McKay, J.L., Factor, S.A., Ting, L.H., 2020. Perception of whole-body motion during balance perturbations is impaired in Parkinson’s disease and is associated with balance impairment. Gait Posture 76, 44–50. 10.1016/j.gaitpost.2019.10.029

Bonini, F., Burle, B., Liégeois-Chauvel, C., Régis, J., Chauvel, P., Vidal, F., 2014. Action Monitoring and Medial Frontal Cortex: Leading Role of Supplementary Motor Area. Science 343, 888– 891. 10.1126/science.1247412

Brown, P., 2003. Oscillatory nature of human basal ganglia activity: relationship to the pathophysiology of Parkinson’s disease. Mov. Disord. 18, 357–363. 10.1002/mds.10358

Cheng, C.-H., Lin, Y.-Y., 2013. Aging-related decline in somatosensory inhibition of the human cerebral cortex. Exp Brain Res 226, 145–152. 10.1007/s00221-013-3420-9

Chong, R.K.Y., Horak, F.B., Woollacott, M.H., 2000. Parkinson’s disease impairs the ability to change set quickly. Journal of the Neurological Sciences 175, 57–70. 10.1016/S0022-510X(00)00277-X

Cromwell, H.C., Mears, R.P., Wan, L., Boutros, N.N., 2008. Sensory Gating: A Translational Effort from Basic to Clinical Science. Clin EEG Neurosci 39, 69–72. 10.1177/155005940803900209

Delorme, A., Makeig, S., 2004. EEGLAB: an open source toolbox for analysis of single-trial EEG dynamics including independent component analysis. J. Neurosci. Methods 134, 9–21. 10.1016/j.jneumeth.2003.10.009

Donoghue, T., Haller, M., Peterson, E.J., Varma, P., Sebastian, P., Gao, R., Noto, T., Lara, A.H., Wallis, J.D., Knight, R.T., Shestyuk, A., Voytek, B., 2020. Parameterizing neural power spectra into periodic and aperiodic components. Nat Neurosci 23, 1655–1665. 10.1038/s41593-020-00744-x

Engel, A.K., Fries, P., 2010. Beta-band oscillations—signalling the status quo? Current Opinion in Neurobiology 20, 156–165. 10.1016/j.conb.2010.02.015

Espenhahn, S., Rossiter, H.E., van Wijk, B.C.M., Redman, N., Rondina, J.M., Diedrichsen, J., Ward, N.S., 2020. Sensorimotor cortex beta oscillations reflect motor skill learning ability after stroke. Brain Communications 2, fcaa161. 10.1093/braincomms/fcaa161

Ghosn, N.J., Palmer, J.A., Borich, M.R., Ting, L.H., Payne, A.M., 2020. Cortical Beta Oscillatory Activity Evoked during Reactive Balance Recovery Scales with Perturbation Difficulty and Individual Balance Ability. Brain Sciences 10, 860. 10.3390/brainsci10110860

Halperin, O., Israeli-Korn, S., Yakubovich, S., Hassin-Baer, S., Zaidel, A., 2020. Self-motion perception in Parkinson’s disease. European Journal of Neuroscience. 10.1111/ejn.14716

Horak, F.B., 2006. Postural orientation and equilibrium: what do we need to know about neural control of balance to prevent falls? Age and Ageing 35, ii7–ii11. 10.1093/ageing/afl077

Horak, F.B., Diener, H.C., Nashner, L.M., 1989. Influence of central set on human postural responses. Journal of Neurophysiology 62, 841–853. 10.1152/jn.1989.62.4.841

Horak, F.B., Macpherson, J.M., 2011. Postural Orientation and Equilibrium, in: Comprehensive Physiology. American Cancer Society, pp. 255–292. 10.1002/cphy.cp120107

Hoseini, N., Sexton, B.M., Kurtz, K., Liu, Y., Block, H.J., 2015. Adaptive Staircase Measurement of Hand Proprioception. PLOS ONE 10, e0135757. 10.1371/journal.pone.0135757

Hussain, S.J., Vollmer, M.K., Iturrate, I., Quentin, R., 2022. Voluntary Motor Command Release Coincides with Restricted Sensorimotor Beta Rhythm Phases. J. Neurosci. 42, 5771–5781. 10.1523/JNEUROSCI.1495-21.2022

Jones, S.R., Kerr, C.E., Wan, Q., Pritchett, D.L., Hämäläinen, M., Moore, C.I., 2010. Cued Spatial Attention Drives Functionally Relevant Modulation of the Mu Rhythm in Primary Somatosensory Cortex. J. Neurosci. 30, 13760–13765. 10.1523/JNEUROSCI.2969-10.2010

Jürgens, U., 1984. The efferent and afferent connections of the supplementary motor area. Brain Research 300, 63–81. 10.1016/0006-8993(84)91341-6

Kilavik, B.E., Zaepffel, M., Brovelli, A., MacKay, W.A., Riehle, A., 2013. The ups and downs of beta oscillations in sensorimotor cortex. Experimental Neurology, Special Issue: Neuronal oscillations in movement disorders 245, 15–26. 10.1016/j.expneurol.2012.09.014

Klug, M., Gramann, K., 2021. Identifying key factors for improving ICA-based decomposition of EEG data in mobile and stationary experiments. European Journal of Neuroscience 54, 8406–8420. 10.1111/ejn.14992

Klug, M., Kloosterman, N.A., 2022. Zapline-plus: a Zapline extension for automatic and adaptive removal of frequency-specific noise artifacts in M/EEG. 10.1101/2021.10.18.464805

Leek, M.R., 2001. Adaptive procedures in psychophysical research. Perception & Psychophysics 63, 1279–1292. 10.3758/BF03194543

Little, S., Bonaiuto, J., Barnes, G., Bestmann, S., 2019. Human motor cortical beta bursts relate to movement planning and response errors. PLOS Biology 17, e3000479. 10.1371/journal.pbio.3000479

Marlin, A., Mochizuki, G., Staines, W.R., McIlroy, W.E., 2014. Localizing evoked cortical activity associated with balance reactions: does the anterior cingulate play a role? Journal of Neurophysiology 111, 2634–2643. 10.1152/jn.00511.2013

Mierau, A., Hülsdünker, T., Strüder, H.K., 2015. Changes in cortical activity associated with adaptive behavior during repeated balance perturbation of unpredictable timing. Front. Behav. Neurosci. 9. 10.3389/fnbeh.2015.00272

Mirdamadi, J.L., Block, H.J., 2020. Somatosensory changes associated with motor skill learning. J. Neurophysiol. 123, 1052–1062. 10.1152/jn.00497.2019

Nachev, P., Kennard, C., Husain, M., 2008. Functional role of the supplementary and pre-supplementary motor areas. Nat Rev Neurosci 9, 856–869. 10.1038/nrn2478

Nguyen, A.T., Hetrick, W.P., O’Donnell, B.F., Brenner, C.A., 2020. Abnormal beta and gamma frequency neural oscillations mediate auditory sensory gating deficit in schizophrenia. Journal of Psychiatric Research 124, 13–21. 10.1016/j.jpsychires.2020.01.014

Oostenveld, R., Oostendorp, T.F., 2002. Validating the boundary element method for forward and inverse EEG computations in the presence of a hole in the skull. Hum Brain Mapp 17, 179–192. 10.1002/hbm.10061

Ostry, D.J., Darainy, M., Mattar, A.A.G., Wong, J., Gribble, P.L., 2010. Somatosensory Plasticity and Motor Learning. Journal of Neuroscience 30, 5384–5393. 10.1523/JNEUROSCI.4571-09.2010

Palmer, J.A., Makeig, S., Kreutz-Delgado, K., Rao, B.D., 2008. Newton method for the ICA mixture model, in: 2008 IEEE International Conference on Acoustics, Speech and Signal Processing. Presented at the ICASSP 2008. IEEE International Conference on Acoustic, Speech and Signal Processes, IEEE, Las Vegas, NV, pp. 1805–1808. 10.1109/ICASSP.2008.4517982

Payne, A.M., Hajcak, G., Ting, L.H., 2019a. Dissociation of muscle and cortical response scaling to balance perturbation acceleration. J. Neurophysiol. 121, 867–880. 10.1152/jn.00237.2018

Payne, A.M., Palmer, J.A., McKay, J.L., Ting, L.H., 2021. Lower Cognitive Set Shifting Ability Is Associated With Stiffer Balance Recovery Behavior and Larger Perturbation-Evoked Cortical Responses in Older Adults. Front Aging Neurosci 13, 742243. 10.3389/fnagi.2021.742243

Payne, A.M., Ting, L.H., 2020a. Worse balance is associated with larger perturbation-evoked cortical responses in healthy young adults. Gait & Posture 80, 324–330. 10.1016/j.gaitpost.2020.06.018

Payne, A.M., Ting, L.H., 2020b. Balance perturbation-evoked cortical N1 responses are larger when stepping and not influenced by motor planning. J Neurophysiol 124, 1875–1884. 10.1152/jn.00341.2020

Payne, A.M., Ting, L.H., Hajcak, G., 2019b. Do sensorimotor perturbations to standing balance elicit an error-related negativity? Psychophysiology 56, e13359. 10.1111/psyp.13359

Pion-Tonachini, L., Kreutz-Delgado, K., Makeig, S., 2019. ICLabel: An automated electroencephalographic independent component classifier, dataset, and website. Neuroimage 198, 181–197. 10.1016/j.neuroimage.2019.05.026

Prins, N., Kingdom, F.A.A., 2018. Applying the Model-Comparison Approach to Test Specific Research Hypotheses in Psychophysical Research Using the Palamedes Toolbox. Frontiers in Psychology 9.

Puntkattalee, M.J., Whitmire, C.J., Macklin, A.S., Stanley, G.B., Ting, L.H., 2016. Directional acuity of whole-body perturbations during standing balance. Gait Posture 48, 77–82. 10.1016/j.gaitpost.2016.04.008

Rossiter, H.E., Davis, E.M., Clark, E.V., Boudrias, M.-H., Ward, N.S., 2014. Beta oscillations reflect changes in motor cortex inhibition in healthy ageing. Neuroimage 91, 360–365. 10.1016/j.neuroimage.2014.01.012

Sawers, A., Hafner, B., 2018. Validation of the Narrowing Beam Walking Test in Lower Limb Prosthesis Users. Arch Phys Med Rehabil 99, 1491–1498.e1. 10.1016/j.apmr.2018.03.012

Shin, H., Law, R., Tsutsui, S., Moore, C.I., Jones, S.R., 2017. The rate of transient beta frequency events predicts behavior across tasks and species. eLife 6, e29086. 10.7554/eLife.29086

Solis-Escalante, T., Stokkermans, M., Cohen, M.X., Weerdesteyn, V., 2020. Cortical responses to whole-body balance perturbations index perturbation magnitude and predict reactive stepping behavior. Eur J Neurosci. 10.1111/ejn.14972

Spooner, R.K., Wiesman, A.I., Proskovec, A.L., Heinrichs-Graham, E., Wilson, T.W., 2019. Rhythmic Spontaneous Activity Mediates the Age-Related Decline in Somatosensory Function. Cerebral Cortex 29, 680–688. 10.1093/cercor/bhx349

Staines, R., McIlroy, W., Brooke, J., 2001. Cortical representation of whole-body movement is modulated by proprioceptive discharge in humans. Experimental Brain Research 138, 235–242. 10.1007/s002210100691

Tanji, J., 1994. The supplementary motor area in the cerebral cortex. Neuroscience Research 19, 251–268. 10.1016/0168-0102(94)90038-8

Taylor, M.M., Creelman, C.D., 1967. PEST: Efficient Estimates on Probability Functions. The Journal of the Acoustical Society of America 41, 782–787. 10.1121/1.1910407

Thibaut, A., Simis, M., Battistella, L.R., Fanciullacci, C., Bertolucci, F., Huerta-Gutierrez, R., Chisari, C., Fregni, F., 2017. Using Brain Oscillations and Corticospinal Excitability to Understand and Predict Post-Stroke Motor Function. Front Neurol 8, 187. 10.3389/fneur.2017.00187

Welch, T.D.J., Ting, L.H., 2008. A feedback model reproduces muscle activity during human postural responses to support-surface translations. J Neurophysiol 99, 1032–1038. 10.1152/jn.01110.2007

Wiesendanger, M., Hummelsheim, H., Bianchetti, M., 1985. Sensory input to the motor fields of the agranular frontal cortex: A comparison of the precentral, supplementary motor and premotor cortex. Behavioural Brain Research 18, 89–94. 10.1016/0166-4328(85)90065-8

Wilson, E.T., Wong, J., Gribble, P.L., 2010. Mapping Proprioception across a 2D Horizontal Workspace. PLoS ONE 5, e11851. 10.1371/journal.pone.0011851

